# Psilocybin exerts distinct effects on resting state networks associated with serotonin and dopamine in mice

**DOI:** 10.1101/751255

**Authors:** Joanes Grandjean, David Buehlmann, Michaela Buerge, Hannes Sigrist, Erich Seifritz, Franz X. Vollenweider, Christopher R. Pryce, Markus Rudin

## Abstract

Hallucinogenic agents have been proposed as potent antidepressants; this includes the serotonin (5-HT) receptor 2A agonist psilocybin. In human subjects, psilocybin alters functional connectivity (FC) within the default-mode network (DMN), a constellation of inter-connected regions that is involved in self-reference and displays altered FC in depressive disorders. In this study we investigated the effects of psilocybin on FC in the analogue of the DMN in mouse, with a view to establishing an experimental animal model to investigate underlying mechanisms. Psilocybin effects were investigated in lightly-anaesthetized mice using resting-state fMRI. Dual-regression analysis identified reduced FC within the ventral striatum in psilocybin-relative to vehicle-treated mice. Refinement of the analysis using spatial references derived from both gene expression maps and viral tracer projection fields revealed two distinct effects of psilocybin: it increased FC between 5-HT-associated networks and elements of the murine DMN, thalamus, and midbrain; it decreased FC within dopamine (DA)-associated striatal networks. These results suggest that interaction between 5-HT- and DA-regulated neural networks contributes to the neural and therefore psychological effects of psilocybin. Furthermore, they highlight how information on molecular expression patterns and structural connectivity can assist in the interpretation of pharmaco-fMRI findings.

## Introduction

Psychiatric disorders are associated with changes in the status of specific neurotransmitter systems, including those of serotonin (5-hydroxytryptamine, 5-HT) and dopamine (DA). Recently, psilocybin, a psychedelic compound derived from “magic mushrooms” with high affinity to the 5-HTA receptor (Nichols, 2004; Halberstadt and Geyer, 2011), has gained in research interest due to its potential to alleviate depression and anxiety (Grob et al., 2011; Griffiths et al., 2016; Carhart-Harris et al., 2017a). Whilst the primary target of its active metabolite psilocin is the 5-HT2A receptor and it binds to a lesser extent to receptor 5-HT1A, psilocybin has also been shown to induce DA release in the nucleus accumbens in rats (Sakashita et al., 2015) and to reduce ^11^C-raclopride binding to the D2 receptor in the caudate-putamen in humans (Vollenweider et al., 1999).

The direct and indirect actions of psilocybin on specific monoamine neurotransmitter systems result in downstream metabolic and physiological effects on the brain. Thus, psilocybin increases global energy metabolism indicated by elevated glucose utilization rates in the frontomedial, frontolateral and anterior cingulate cortices as well as in basal ganglia (Vollenweider et al., 1997). Also, somewhat counterintuitively, reduced cerebral blood flow (CBF) in several brain regions including cingulate cortex, thalamus and striatum, has been reported (Carhart-Harris et al., 2012). In blood oxygen level-dependent functional magnetic resonance imaging (BOLD fMRI) studies, psilocybin decreased functional connectivity (FC) within the default-mode network (DMN), and between the anterior DMN and task-positive networks (Carhart-Harris et al., 2012; Carhart-Harris et al., 2013; Roseman et al., 2014). Psilocybin therefore impacts on FC in various networks throughout the human brain, with the main effect centred on the DMN. This is consistent with the concept that psilocybin’s psychoactive effects are mediated by networks underlying self-referential processing (Carhart-Harris et al., 2014). The mechanisms, whereby psilocybin acts on functional activity within the DMN and other resting-state networks (RSNs), remain poorly understood, including how the 5-HT system is involved.

The contributions of the monoamine systems 5-HT and DA to FC are challenging to understand. The midbrain nuclei hosting 5-HT (raphe nuclei) and DA (ventral tegmental area, substantia nigra pars compacta) cell bodies are not included in canonical RSNs (Damoiseaux et al., 2006). Using BOLD fMRI, correlational analysis of signal traces extracted from midbrain nuclei typically fails to capture evidence of functional coupling to distal regions. Integrating complementary information with FC analysis might overcome this limitation and reveal contributions of midbrain nuclei to FC. In mouse models, integration of whole-brain imaging with region-specific gene expression (Lein et al., 2007; Ng et al., 2009; Hawrylycz et al., 2012) and axonal projection maps (Oh et al., 2014) derived from the Allen Institute for Brain Science (AIBS) offers the opportunity to investigate the molecular neuropharmacology of psychoactive agents in spatio-temporal dimensions.

Here we hypothesized that i) acute psilocybin administration in mice increases FC within and between elements of the rodent DMN (Sforazzini et al., 2014; Stafford et al., 2014; Gozzi and Schwarz, 2016), and ii) gene expression and axonal projection maps related to specific monoamine systems can be used to identify specific contributions of these systems to the psilocybin-induced changes in FC pattern observed. To test these hypotheses, we acquired resting-state fMRI data from mice treated with psilocybin or vehicle. Conventional data analysis did not reveal any drug-induced alterations within elements of the rodent DMN. Rather, we observed a significant decrease in striatal FC, pronounced in the ventral striatum, following psilocybin. Gene expression maps for the 5-HT2A receptor or the D2 receptor, and axonal projection field maps for dorsal raphe nucleus (DRN) 5-HT neurons and ventral tegmental area (VTA) DA neurons, were used as spatial references for identifying 5-HT and DA contributions to the RSN changes induced by psilocybin. This novel iterative approach revealed psilocybin effects not evident from conventional fMRI analysis, and demonstrated that psilocybin does act on FC within the rodent DMN, albeit to a lesser extent compared to its FC effects in the ventral striatum. These findings provide new translational insights into the neuropharmacology of psilocybin.

## Material and methods

### Animal preparation and psilocybin administration

This study was conducted in accordance with federal guidelines and under a license from the Zürich Cantonal Veterinary Office (149/2015). A total of 50 young-adult male C57BL/6 mice (Janvier, Le Genest-St Isle, France) weighing 25 ± 1.2 g were studied. Animals were maintained in standard housing with food and water available *ad libitum* and in a reversed 12:12 day/night cycle (light off: 07:00-19:00 h). Anaesthesia was induced with isoflurane 3.5% in a mixture of 20% O2 and 80% air. Isoflurane was reduced to 2%, animals were then intubated endotracheally and ventilated mechanically at 80 breaths/min, positioned on an MR-compatible cradle, and a cannula was positioned in the tail vein. Psilocybin was dissolved in sterile water with tartaric acid and infused i.v. during 5 min with either 1 mg/kg (n = 13) or 2 mg/kg (n = 12) or vehicle only (n = 15). These doses were chosen because they were demonstrated previously to elicit hallucinogen-specific behaviour in mice (Gonzalez-Maeso et al., 2007). In contrast, doses comparable to those used in humans failed to elicit detectable hemodynamic changes in the resting BOLD signal in rats (Carhart-Harris et al., 2012; Spain et al., 2015). Following the infusion, a bolus of medetomidine (0.05 mg/kg, Domitor, medetomidine hydrochloride; Pfizer Pharmaceuticals, Sandwich, UK) together with pancuronium bromide (0.2 mg/kg, Sigma-Aldrich, Steinheim, Germany) was injected i.v. and isoflurane was reduced to 1.5%. After 5 min, isoflurane was further reduced to 0.5% and medetomidine (0.1 mg/kg/h) and pancuronium bromide (0.4 mg/kg/h) were infused during the remainder of the imaging acquisition protocol as in (Grandjean et al., 2014). Functional imaging began 20 min after completion of the psilocybin or vehicle infusion. To test for potential effects of psilocybin on cardiovascular function that would lead to confounding effects on brain FC, in a separate group of mice (n = 10) electrocardiogram recordings were made whilst otherwise applying the same protocol as for the MRI experiment. Electrocardiogram recordings were performed using 3 sets of electrodes (SA instruments Inc, Stony Brook, NY, USA) inserted into the front paws and left hind paw. All animals in the study were included in the data analysis.

### Image acquisition

MRI experiments were conducted using a Bruker Biospec 94/30 small animal MR system (Bruker BioSpin MRI, Ettlingen, Germany) operating at 400 MHz (9.4 T). A linearly polarized volume resonator was used for transmission, together with a four-element receiver-only cryogenic phased array coil (Bruker BioSpin AG, Fällanden, Switzerland). A multi-echo gradient-echo echo planar imaging (ME-EPI) sequence was used to acquire BOLD fMRI with: repetition time 1500 ms, echo time [11, 17, 23] ms, flip angle 60°, matrix size 60×30, field of view 18.2×9 mm^2^, number of coronal slices 20, slice thickness 0.3 mm, slice gap 0.05 mm, 800 volumes, acceleration factor 1.4, horizontal field of view saturation slice to mask the lower portion of the mouse head, and 250000 Hz bandwidth. Images were pre-processed in AFNI (AFNI_16.1.26, https://afni.nimh.nih.gov/), using the meica.py script (Kundu et al., 2012) to reconstruct the ME-EPI and remove non-BOLD noise contributions. Reconstructed ME-EPIs were transformed to a study EPI template in ANTS (V2.1, http://picsl.upenn.edu/software/ants/) using linear affine and nonlinear SyN diffeomorphic transformation (Avants et al., 2011).

### Experimental design and statistical analysis

The complete dataset used in this study is fully available online (PROJECT_ID: Mouse_rest_psilocybin, https://openneuro.org/datasets/ds001725)

Acquisition was carried out in two separate subsets with respect to the experimenter (JG or DB): subset 1 comprised vehicle = 8, 1 mg/kg n = 6, 2 mg/kg n = 5 mice, and subset 2 comprised vehicle n = 7, 1 mg/kg n = 7, 2 mg/kg n = 7) mice. Voxel-based within-network functional connectivity analysis was carried out using a dual-regression framework (Filippini et al., 2009). Briefly, spatially delineated reference maps are used to extract an averaged BOLD time course for each scan. A general linear model was used to regress these time courses into the individual scans, to obtain parameter estimate maps indicative of network strength at every voxel for each corresponding reference map. The spatial reference maps used in the dual-regression analyses were either 17 references RSNs as determined with group independent component analysis (ICA) in (Zerbi et al., 2015), *in-situ* hybridization (ISH) gene expression maps (Lein et al., 2007), or viral tracer maps (Oh et al., 2014). To confirm the spatial distribution of the ventral striatum network, seed-based maps were obtained by extracting time courses from a selected seed region in the ventral striatum, corresponding to the peak psilocybin effect in the dual-regression analysis. The BOLD signal from the seed regressing them in a general linear model into individual scans from the vehicle group to obtain parameter estimate maps denoting FC relative to the seed region. Group-level statistical analysis was carried out with a non-parametric permutation test (5000 permutations), with parameter estimate maps from mice administered with either psilocybin 1 mg/kg or 2 mg/kg compared to vehicle, or a combined psilocybin group (1 and 2 mg/kg) compared to vehicle. The statistical correction for multiple hypothesis testing was carried out in two stages: firstly using threshold-free cluster enhancement (TFCE) and second using Bonferroni correction to account for the number of reference maps investigated. Between-network analysis was carried out within the FSLNets framework using false discovery rate (FDR) correction (FMRIB Software Library v5.0, fsl.fmrib.ox.ac.uk). Parametric one-sample *t*-tests were carried out in the seed-based analysis. Statistical maps are shown as colour-coded *t*-statistic overlay on the AMBMC template (www.imaging.org.au/AMBMC/AMBMC) with a threshold set at p<0.05 (TFCE and Bonferroni corrected). Region-of-interest (ROI) analysis was performed using a parcellation based on the AIBS atlas. Statistical analysis at the ROI level was performed with a parametric linear model with contrast analysis and corrected with FDR. Descriptive statistics are given as mean ± 1 standard deviation. Estimation statistics are provided as mean differences ± 95^th^ confidence interval estimated by bootstrap resampling (https://www.estimationstats.com) (Ho et al., 2019).

### AIBS database search

Maps of gene expression and projection field were obtained from the AIBS database (http://mouse.brain-map.org/). ISH gene expression maps relating to genes encoding 5-HT and DA receptors were searched using “Htr” and “Drd” terms, respectively. Expression maps were selected based on the presence of detectable expression and absence of artefact in the reconstructed images. The maps from the following experiments were downloaded using the AIBS application programming interface: Htr1a 79556616, Htr1b 583, Htr1f 69859867, Htr2a 81671344, Htr2c 73636098, Drd1 71307280, Drd2 81790728. Viral tracer maps were obtained by selecting AIBS experiments carried out in lines expressing the Cre recombinase in 5-HT and DA neurons for the DRN (114155190) and VTA (160539283), respectively. Maps were spatially transformed to reference space using ANTS and normalized to a [0, 100] range. Injection site and white matter fibres were masked from tracer injection maps using white matter masks. An anatomic gene expression atlas (AGEA) correlation map denoting the correlation between the gene expression profile obtained from a seed region in the ventral striatum and that of the gene expression profiles across every remaining voxel in the brain was obtained from the AIBS database. In effect, an AGEA map indicates the degree of resemblance in gene expression across the whole brain relative to a target region. The correlations were established on the basis of 4376 ISH maps previously selected for genes expressed in the brain (Lein et al., 2007). A Spearman’s rank correlation was used to compare the spatial overlap of the AGEA map to the seed-based map estimated with a seed in the ventral striatum. Additional comparisons between ISH maps and the 17 references RSNs were also performed with Spearman’s rank correlations. The significance of a correlation between *Drd2* or *Htr2a* gene expression maps and individual RSN maps was determined as the 95^th^ percentile of a null distribution estimated using the remaining 4374 ISH gene expression maps (the original 4376 ISH maps minus those for *Drd2* and *Htr2a*).

## Results

### Psilocybin predominantly affects functional connectivity within the ventral striatum

To test for potential confounding cardiovascular-mediated effects of psilocybin on brain FC, heart rate was monitored in a separate group of mice using electrocardiogram recordings and applying the same protocol as for the MRI experiment. Heart rate was 302 ± 30 beats per minute (bpm) in mice (n = 5) that received2 mg/kg psilocybin and 296 ± 27 bpm in mice (n = 5) that received vehicle; psilocybin was without a significant effect on heart rate therefore (two-sample *t*-test, *t* = 0.32, p-value = 0.75), consistent with observations in humans (Hasler et al., 2004).

Using the 17 RSNs established with ICA in a previous study (Zerbi et al., 2015), dual-regression analysis identified that psilocybin led to reduced FC within the ventral striatum network of approximately 37%, with the effect similar at the two psilocybin doses (Figure 1, mean difference _1 mg/kg minus Veh_ = −15.21 ± [−21.67, −8.22], mean difference _2 mg/kg minus Veh_ = −13.35 ± [−20.85, −5.16]). As for the ventral striatum, also for other RSNs there was no significant dose-dependent psilocybin effect, and therefore the two dose groups were pooled to enhance statistical power and the analysis run again. The psilocybin effect of reduced FC in the ventral striatum was robust and reproducible, as the two subsets revealed similar results (Figure S1). Psilocybin effects on FC were obtained within three additional RSNs, the piriform cortex, dorsal striatum and lateral striatum (Figure S2); however, these effects did not survive the second stage of statistical correction, namely Bonferroni threshold. In a network analysis, there were no significant psilocybin effects on between-network FCs for any pair of RSNs after full correction. We conclude that the striatal networks, and in particular the ventral striatum RSN, were the most responsive to psilocybin, therefore.

**Figure 1.**
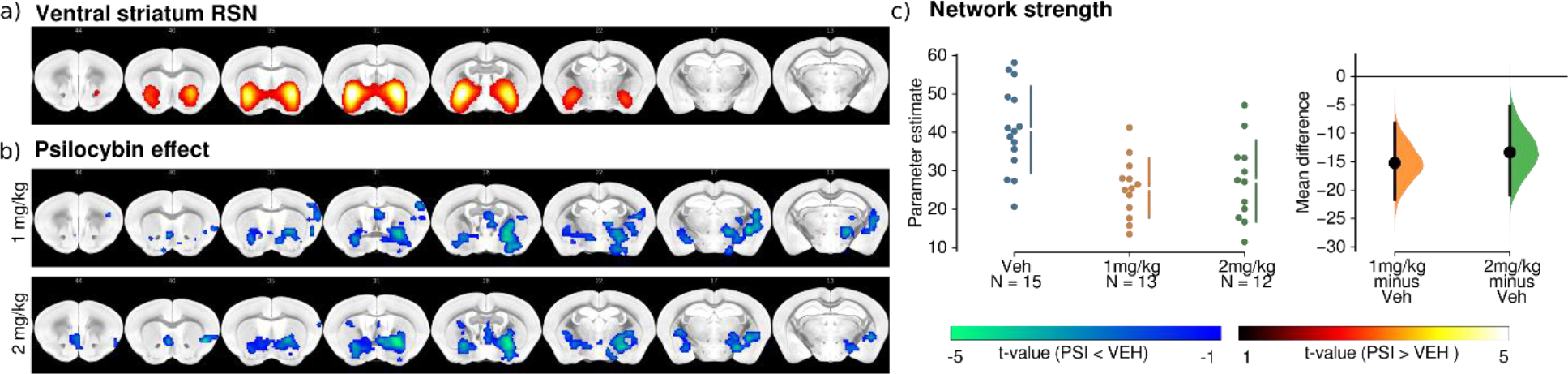
a) Reference map for the ventral striatum RSN and (b) statistical non-parametric maps estimated with dual-regression indicating decreased FC within the striatal RSN and extending to the pallidum in the psilocybin groups (1 mg/kg, 2 mg/kg) relative to vehicle. c) Network strength extracted from the ventral striatum in the 3 groups. A comparable mean difference was found in both psilocybin groups relative to the control group. Reference RSN is shown as colour-coded overlay. Statistical non-parametric maps are shown as colour-coded t-statistics (p<0.05 corrected with TFCE and Bonferroni correction). Mean differences are plotted as bootstrap sampling distributions. Black dots indicate mean differences. Vertical error bars indicate 95^th^ confidence intervals. Vehicle n = 15, psilocybin 1 mg/kg n = 13, psilocybin 2 mg/kg n = 12.

### Ventral striatum RSN overlaps with a gene cluster characterized by high DA receptor gene expression

Recent work has demonstrated a tight link between resting-state fMRI RSNs and spatial patterns of gene expression (Richiardi et al., 2015). To investigate gene expression in the ventral striatum (the RSN that was most sensitive to psilocybin), the highest t-value voxel in the statistical map (Figure 1) denoting the peak location of the psilocybin effect, was used as a reference region for a seed-based analysis in the vehicle group (Figure 2). The resultant seed-based maps for ventral striatum FC were consistent with the spatial delineation of the corresponding reference RSN in the ICA atlas established for untreated mice (Zerbi et al., 2015). The gene expression profiles of 4376 genes in brain tissues were extracted from the same seed region as above, and compared to those of the remaining voxels in the brain. This determined a cluster of voxels co-expressing the same genes. The gene expression correlation map strongly overlapped with the seed-based map for these ventral striatum voxels (rho = 0.356, p < 10^−16^), indicative of a conserved topology between gene expression and intrinsic functional activity. This gene expression cluster was enriched with, among other genes, those encoding for the D1 and D2 receptors (1.75 and 2.06 fold increase in *Drd1* and *Drd2* expression, respectively, compared with voxels outside the cluster). This indicates that the RSN predominantly affected by psilocybin is enriched in DA receptors.

**Figure 2.**
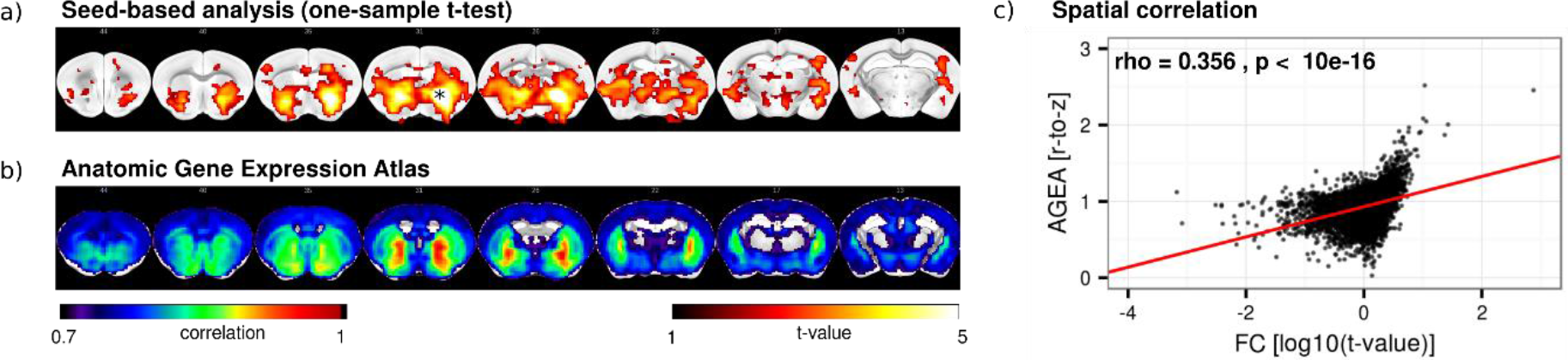
a) One-sample t-test maps performed in vehicle-treated mice (n = 15) for seed-based FC maps estimated with a 2×2 voxel seed region placed in the ventral striatum (asterisk), which was found to display the highest level of significance regarding psilocybin effects (see Figure 1). FC relative to the seed co-localizes with the ventral striatum in the vehicle group. b) The gene co-expression cluster obtained from the anatomic gene expression atlas (AGEA) for the same seed indicates voxels with correlating gene expression patterns to the seed voxel. c) Its spatial extent overlaps with the FC cluster. Voxel-wise comparison across all brain voxels resulted in a strong correlation between the two maps (rho=0.356, p < 10^−16^). Seed-based FC maps are shown as t-statistics (p<0.05 uncorrected), Gene co-expression is shown as colour-coded correlation coefficient relative to the seed.

### 5-HT2A and D2 receptor expressions are localized in different RSNs

The previous analysis identified that the ventral striatum network overlapped with a gene co-expression cluster enriched in DA receptors. Next, the analysis was extended to investigate for inter-relationships between the remaining RSNs and the gene expression maps for 5-HT2A (*Htr2a*) and D2 (*Drd2*) receptors. D2 was selected because psilocybin decreased the binding potential of the D2 PET ligand ^11^C-raclopride indicating increased receptor occupancy by endogenous dopamine (Vollenweider et al., 1999), and 5-HT2A is the main target of psilocin (Nichols, 2004; Halberstadt and Geyer, 2011) and necessary for its hallucinogenic effects (Vollenweider et al., 1998; Gonzalez-Maeso et al., 2007). To test for spatial associations of RSNs with either *Drd2* or *Htr2a* expression, Spearman rank correlation was calculated for each of the 17 RSNs (Figure 3a) and the gene expression maps extracted from the AIBS database (Figure 3b and c, Figure S3a). Significance was assessed relative to a null distribution generated from the Spearman rank correlations between each RSN and the gene expression maps for the other 4374 genes expressed in brain tissues (Figure 3d). The *Drd2* gene expression map was significantly spatially correlated with the dorsal, lateral, and ventral striatum networks (rho = [0.47, 0.34, 0.41], p-value = [0.004, 0.008, 0.001], respectively). For the *Htr2a* gene expression map, significant spatial correlations were observed with the piriform cortex, prefrontal cortex, and ventral striatum networks (rho = [0.29, 0.22, 0.20], p-value = [0.03, 0.03, 0.04], respectively). We conclude that *Htr2a* and *Drd2* are expressed within these distinct RSNs. Interestingly, the ventral striatum network, presenting the strongest within-network psilocybin effect, was found to overlap with both receptor expression profiles.

**Figure 3.**
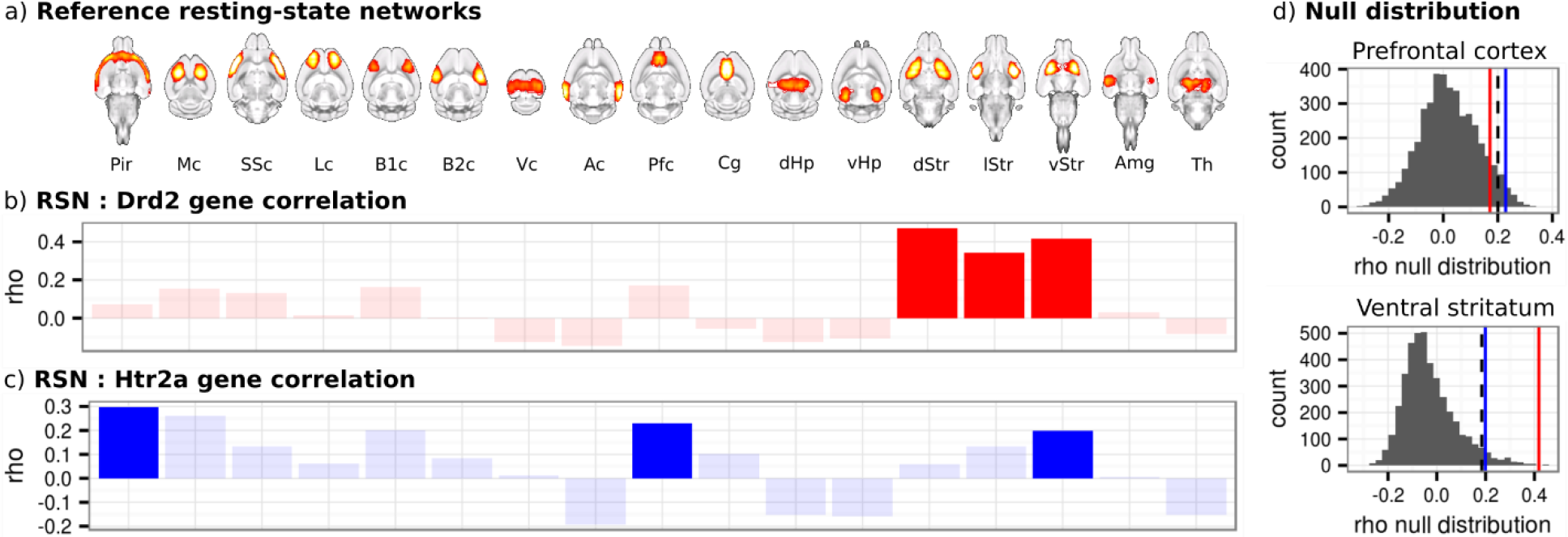
a) The 17 reference mouse brain RSNs b) Correlation between Drd2 gene expression map and the RSNs indicate significant correlations for the dorsal, ventral, and lateral striatum networks (dStr, lStr, vStr). Correlations below the significance threshold are shown in light shades. c) Correlation between Htr2a gene expression map and the RSNs indicates significant correlations in the piriform (Pir) and prefrontal (Pfc) cortex networks and the vStr network. d) A null distribution denoting the spatial correlation between 4374 gene expression and the prefrontal cortex or ventral striatum are represented as grey histograms. These report the distribution of the correlation coefficients between the RSN spatial map and a sample of gene expression maps selected for their expression in brain tissues. The dashed lines indicate the 95^th^ percentile of that null distribution used as a significance threshold, red lines indicate value for correlation to Drd2 gene expression map, and blue lines to Htr2a gene expression map.

### Using 5-HT and DA gene expression and projection field to increase acuity of analysis of psilocybin effects on resting-state FC

To differentiate the effects of psilocybin on 5-HT- and DA-associated networks across the whole brain, a dual-regression analysis was carried with brain-wide *Drd2* and *Htr2a* gene expressions used as spatial reference maps (Figure 4a and b, Figure S3a). The averaged resting BOLD time series from the regions delineated in these reference maps were extracted and used to establish FC between these reference regions and the remaining voxels across the whole-brain. The dual-regression analysis revealed reduced FC between the *Drd2* expressing regions and the striatum, pallidum and thalamus in psilocybin versus vehicle-treated mice, comparable to the results obtained previously (Figure 1). A weak opposite effect, i.e. increased FC due to psilocybin, was found between *Htr2a* expressing regions and the left dorsal striatum, although this effect did not survive statistical correction in the subsequent ROI analysis (Figure 4b). Analyses carried out for other 5-HT receptor gene-expression maps did not reveal significant clusters. To confirm these results, we used viral tracer maps, obtained by virus injection targeting DA neurons in the VTA or 5-HT neurons in the DRN, as spatial references in the dual-regression analysis (Figure 4c and d, Figure S3b). In agreement with the results based on gene expression levels, psilocybin induced a significant decrease in FC between the VTA projection fields and the medial and posterior subfields of the striatum. In contrast, psilocybin administration increased FC between the DRN projection fields and the dorsal striatum, retrosplenial cortex, visual cortex, thalamus and midbrain. We conclude that psilocybin administration exerts differential effects on 5-HT- and DA-associated networks; namely, it increases FC between the 5-HT system and several striatal, cortical, and thalamic regions, and decreases FC between the DA system and striatal regions. It is noteworthy that the 5-HT-related effects were not detected when applying conventional RSN analysis, i.e. examining within-network and between-network FC relative to ICA-based reference mouse RSNs. Hence, the inclusion of prior information concerning the origins and projection pathways of specific neurotransmitter systems, based on gene expression patterns and/or structural connectivity derived from viral tracer maps, can enhance the sensitivity of FC analysis.

**Figure 4.**
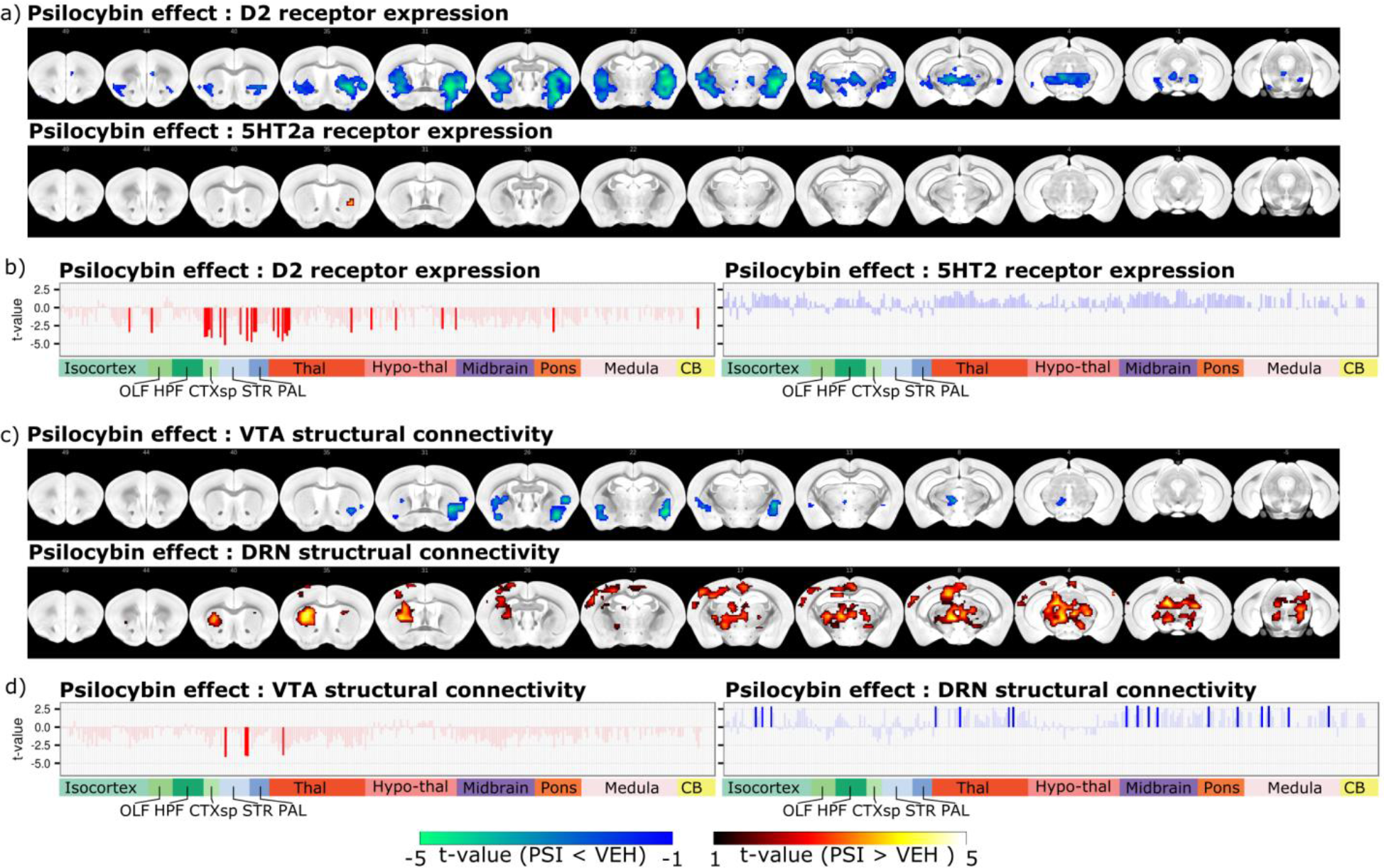
a) Statistical non-parametric maps showing psilocybin effect relative to vehicle for dual-regression analysis carried out with gene expression maps as spatial references. FC was reduced between Drd2 gene expressing regions and in the striatum, pallidum, and thalamus in the psilocybin groups relative to vehicle. A small cluster was present in the striatum denoting an increase in FC between Htr2a expressing regions and this striatum in the psilocybin groups relative to vehicle. b) ROI analysis confirms the presence of significant effects of psilocybin in ROIs found in the striatum, pallidum and thalamus relative to Drd2 expressing regions, denoting decreased FC in the psilocybin mice relative to controls. There was no significant effect relative to Htr2a expression regions. c) Statistical non-parametric maps obtained using the VTA and DRN projection fields as reference maps. Reduced FC is observed between the striatum and pallidum, and regions receiving VTA projection, in the psilocybin groups relative to vehicle. Increased FC is observed between the dorsal striatum, retrosplenial cortex, visual cortex, thalamus, and midbrain, and regions receiving DRN projections, in the psilocybin groups relative to vehicle. d) ROI analysis confirms the presence of significant effects relative to VTA and DRN projection fields. Statistical non-parametric maps are shown as colour-coded t-statistics (p<0.05 corrected with FDR and Bonferroni correction). Bar plots indicate t-statistics for parameter estimate comparisons extracted using AIBS Atlas ROIs. Values below the significance threshold (FDR corrected) are shown in light shades. Colour labels indicate ontological structures: OLF=Olfactory areas, HPF=Hippocampal formation, CTXsp=Cortical sub-plate, STR=Striatum, PAL=Pallidum, Thal=Thalamus, Hypo-thal=Hypothalamus, CB=Cerebellum. Reference spatial maps used to delineate Drd2- and Htr2a-expressing regions, as well as VTA and DRN projections are indicated in **Figure S3.**

## Discussion

We hypothesized that psilocybin administration would decrease FC in the murine DMN-, similar to the effects observed in the human DMN (Carhart-Harris et al., 2012; Carhart-Harris et al., 2013). However, this was not apparent in the initial analysis. Instead, psilocybin reduced FC in the ventral striatum. According to gene expression maps, the RSN ventral striatum is, enriched in DA receptors and, to a lesser extent, 5-HT2A receptors. We then devised a novel analytical approach based on pre-defined projection fields derived from gene expression patterns and viral tracing maps to evaluate specific associations of 5-HT and DA-associated regions with estimated resting-state FC in the corresponding RSNs. This identified distinct effects of psilocybin on RSNs enriched in these two monoamine systems; namely, psilocybin increased FC between 5-HT-associated regions and RSNs of the mouse DMN, and decreased FC between the DA associated regions and striatal RSNs.

The DMN is considered to be one of the most important systems underlying higher cognitive functions (Raichle et al., 2001), and within- and between-region FC is increased in depression (Sheline et al., 2010) and animal models thereof (Grandjean et al., 2016). In human volunteers receiving either psilocybin or placebo, Carhart-Harris and colleagues reported psilocybin-induced: reduction of FC between the DMN regions prefrontal cortex and posterior cingulate cortex (Carhart-Harris et al., 2012); increase in FC between the DMN and task-positive networks (Carhart-Harris et al., 2013); and a general increase in between-networks FC (Roseman et al., 2014). In patients with treatment-resistant depression, psilocybin was shown to normalize DMN activity (Carhart-Harris et al., 2017b). These studies indicate that the DMN is the main target for psilocybin-mediated effects. In mice, by contrast, psilocybin effects on the DMN RSNs were minor compared to the effects on the ventral striatum. This apparent discrepancy could be in part due to species-specific differences in psilocybin pharmacology and methodologies. Firstly, in a rat study, Spain et al. (2015) estimated that a dose of 0.03 mg/kg psilocin, psilocybin’s active metabolite, corresponds to the 2 mg psilocybin dose administered in the human studies carried by Carhart-Harris and colleagues (Spain et al., 2015). However, they did not observe a drug-induced change in the BOLD signal at this dose. In a pilot study using 0.5 mg/kg psilocybin in mice, we only detected weak effects on mouse striatal RSN FC recapitulating the results presented above (Figure S4). At a higher psilocin dose of 2 mg/kg, Spain et al. observed reduced BOLD signal relative to baseline in the cingulate and retrosplenial cortical regions and increased BOLD signal in the amygdala and hypothalamus (Spain et al., 2015). Our present findings agree in part with these: we observed increased FC between regions expressing *Htr2a* and the cingulate and retrosplenial cortices; however, we did not observe effects within the amygdala or hypothalamus. Both of these latter regions are located ventrally in the brain and are therefore susceptible to exposure to magnetic susceptibility gradients leading to geometric distortion and BOLD signal loss (intravoxel dephasing). This can render signal detection somewhat variable in these regions and may account for the discrepancies. Second, in contrast to human studies, the rodent fMRI studies conducted with psilocybin to-date have used anaesthesia. Even though light anaesthesia was used here, interaction with the action of psilocybin cannot be excluded. This could be particularly the case for higher-order cortical networks, such as the mouse analogue of the DMN, which reportedly are affected by under light anaesthesia (Deshpande et al., 2010; Liu et al., 2015).

Psilocin, the active metabolite of psilocybin, is a 5-HT analogue and has been shown to bind to 5-HT2A and 5-HT1A receptors, and with lesser affinity to other 5-HT receptors (Nichols, 2004; Halberstadt and Geyer, 2011). The expression of 5-HT2A receptors on pyramidal glutamate neurons in the cortex was demonstrated as necessary and sufficient for induction of hallucinogen-specific behavioural responses (head twitch, ear scratch) in mice following i.v. administration of 5-HT2A agonists including psilocybin (Gonzalez-Maeso et al., 2007). In human, psilocybin effects on the DA system have been demonstrated by PET study: psilocybin led to decreased D2 receptor occupancy by ^11^C-raclopride, consistent with an increase in endogenous dopamine, in the dorsal striatum (Vollenweider et al., 1999). Using microdialysis, psilocin was shown to increase DA release in the ventral striatum of rats, whereas 5-HT release in this region was not affected. However, psilocybin did increase 5-HT release in the prefrontal cortex (Sakashita et al., 2015). These observations are in line with the psilocybin-induced reduced FC within DA-associated regions observed in our study and suggest a potential mechanism for the reduced FC within the striatum, namely increased DA release in this region. Importantly, the analysis framework developed in this study, using either structural projections or gene expression data concurrently with fMRI analyses, may allow for the overcoming of apparent limitations imposed by the generic physiological nature of the resting-state BOLD signal. Associating distinct FC patterns to either gene expression or structural connectivity priors facilitates the identification of receptor-specific or nucleus-specific contributions to FC. These analytical methods are complementary to those that apply optogenetics or chemogenetics to regulate monoamine release and study effects thereon of brain-wide hemodynamic responses (Giorgi et al., 2017; Grandjean et al., 2019).

We are aware of potential limitations associated with our approach. Firstly, psilocin administration has been associated with decoupling between the hemodynamic response and neuronal activity in rats during short duration, high frequency (>10 Hz) whisker sensory stimulation (Spain et al., 2015); however, this was not the case for lower frequency paradigms. Moreover, this effect would not be expected to be confined to specific RSNs, hence the relevance of this observation in the context of resting-state FC remains moot. Second, database resources of gene expression or viral tracer maps differ in map coverage and quality. Image artefact or misregistration could impact the analysis when used as priors. For instance, only a subset of the 5-HT gene expression maps available on the AIBS database met our quality assurance criterion, restricting the scope of our analysis to this subset of 5-HT genes. Finally, we have used a relatively high dose of psilocybin compared to studies in humans (Carhart-Harris et al., 2012). Species-specific pharmacokinetic and pharmacodynamic properties may account for the large difference in drug dose required to elicit pharmacological responses in humans and rodents.

In conclusion, we have demonstrated robust effects of psilocybin on FC within the striatal RSNs. Incorporation of neurotransmitter-specific gene expression and viral tracer maps into our analysis enabled further refinement. In particular, we found differential psilocybin effects in 5-HT versus DA associated networks: increased FC in elements of the rodent DMN, thalamus, and midbrain attributed to 5-HT versus decreased FC in the striatum, a DA projection area rich in D2 receptors. While the implication of DMN nodes in our analysis supports our original hypothesis, this effect was comparatively small in comparison to the effect on FC within the ventral striatum. Finally, we demonstrate the application of a novel analytical approach to refine the interpretation of FC changes in RSNs. Including information on gene expression and viral tracer maps, available from large neuroscience databases, in the analysis of neuroimaging data, allows for the annotation of FC data with a molecular signature, and thereby provides mechanistic insight into factors involved in normal and pathological network functioning.

