## Supplemental Material for "Psilocybin exerts distinct effects on resting state networks associated with serotonin and dopamine in mice"

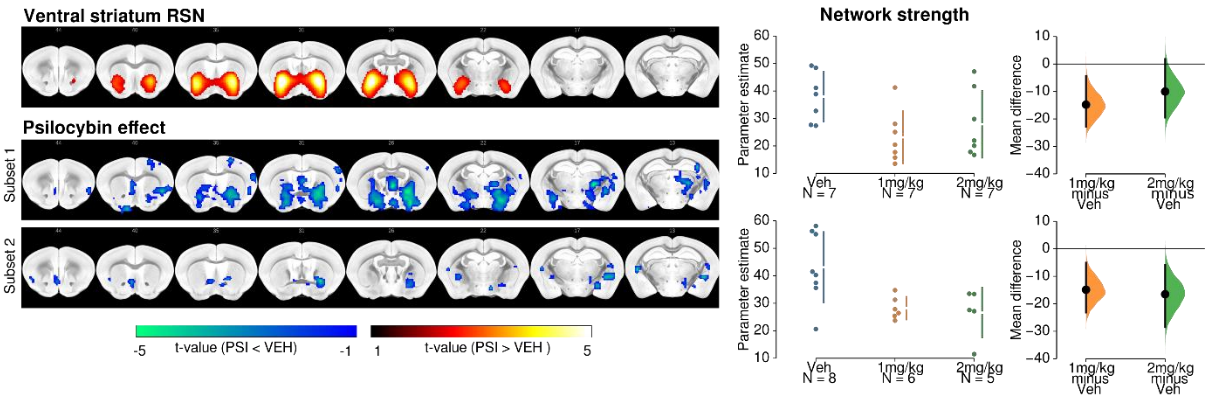


**Figure S1 |** Reproducibility of the psilocybin effect in two independent sets of acquisition. Combined dose psilocybin effect was observed in comparable voxels in subset 1 (vehicle n = 7, psilocybin 1 mg/kg n = 7, psilocybin 2 mg/kg n = 7) and subset 2 (vehicle n = 8, psilocybin 1 mg/kg n = 6, psilocybin 2 mg/kg n = 5). Reference RSN is shown as colour-coded overlay, Statistical non-parametric maps are shown as colour-coded t-statistics ($p<0.05$ corrected with TFCE and Bonferroni correction). Mean differences are plotted as bootstrap sampling distributions. Black dots indicate mean differences. Vertical error bars indicate 95th confidence intervals.

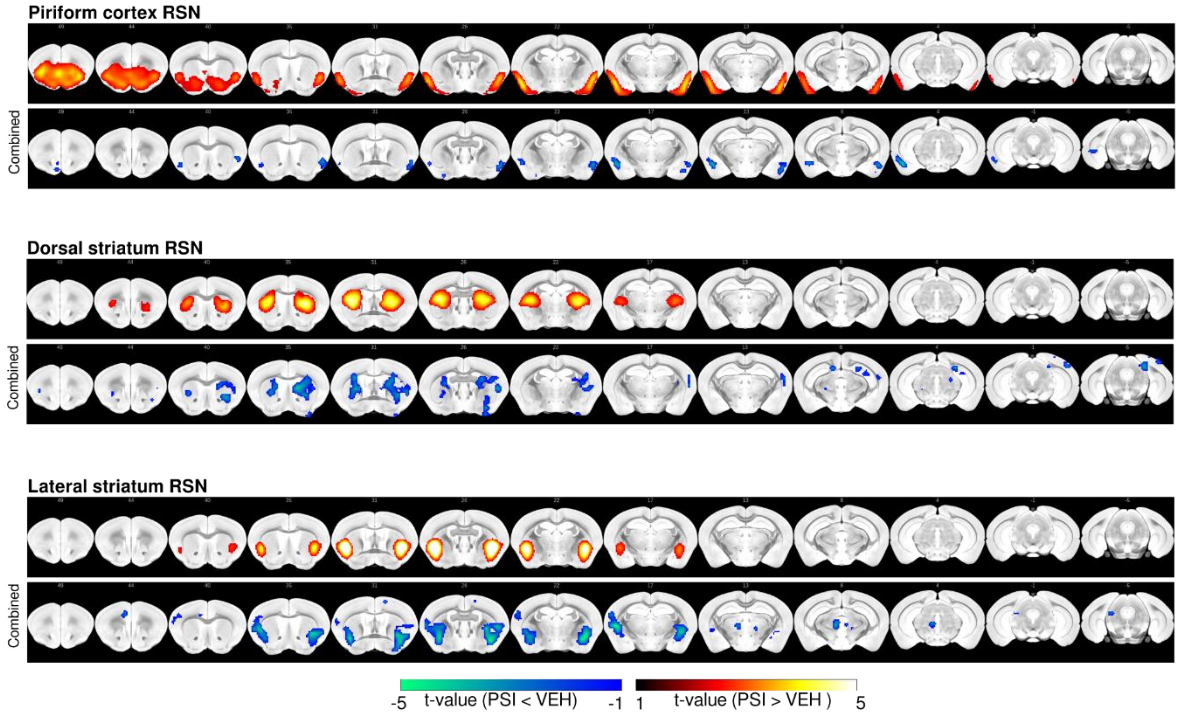


**Figure S2 |** Psilocybin effect in piriform cortical, dorsal striatum, and ventral striatum resting-state networks (RSNs). Reference networks are displayed as colour-coded overlays (upper row in each panel). Psilocybin administration led to a reduction of within-network FC in the piriform, dorsal striatum and lateral striatum networks; however, these effects did not survive Bonferroni correction. Statistical non-parametric maps are shown as colour-coded t-statistics ($p<0.05$ corrected with TFCE). Psilocybin groups (1 and 2 mg/kg) were combined for the analysis.

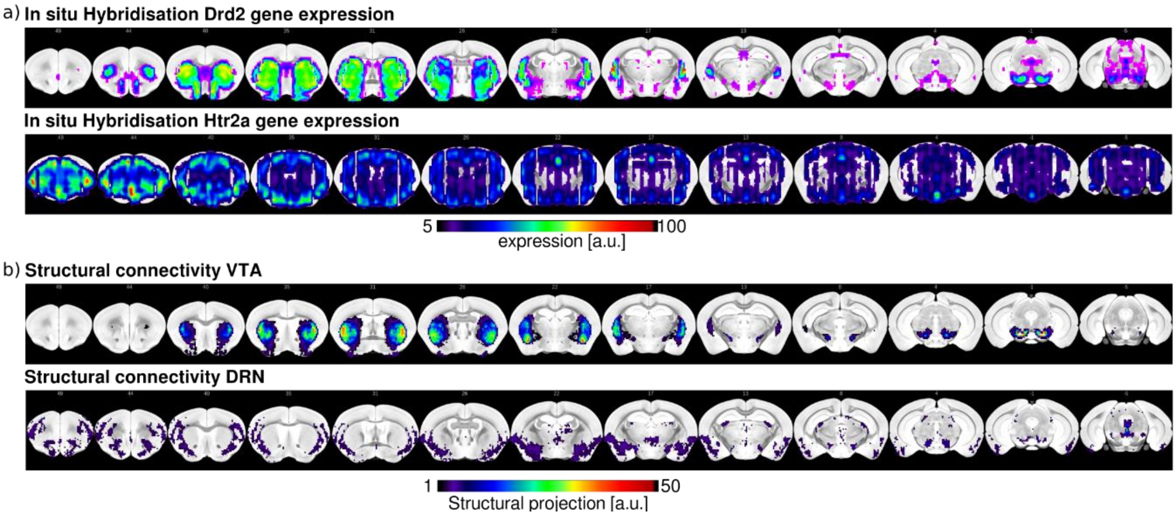


Figure S3 | a) Reference gene expression and (b) structural connectivity maps obtained from the Allen Institute for Brain Sciences database. Specifically, D2 receptor gene Drd2 expression map was obtained from experiment 81790728 and 5-HT2A receptor gene Htr2a map from 81671344. The VTA projection map was obtained from experiment 160539283, which used a Slc6a3-Cre mouse line expressing Cre recombinase in dopaminergic neurons. The DRN projection map was obtained from experiment 114155190, which used a Slc6a4-Cre mouse line expressing Cre recombinase in 5-HT neurons.

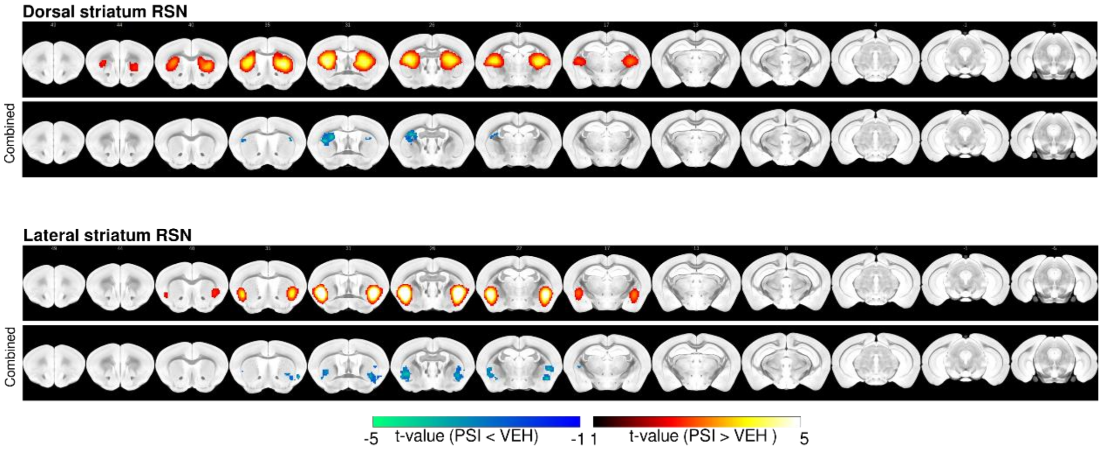


**Figure S4 |** Psilocybin effect in dorsal striatum and lateral striatum resting-state networks (RSNs) in a pilot experiment. Both vehicle and psilocybin (0.5 mg/kg) scans were acquired sequentially in N=12 animals during the same session. Vehicle was administered i.v. 20 min prior to a first fMRI acquisition, and then psilocybin was administered i.v., and a second fMRI scan was acquired 20 min later. Reference networks are displayed as colour-coded overlays (upper row in each panel). Psilocybin administration led to a reduction of within-network FC in the dorsal striatum and lateral striatum networks; however, the effect did not survive Bonferroni correction. Statistical non-parametric maps are shown as colour-coded t-statistics ($p<0.05$ corrected with TFCE). The complete dataset used in this pilot is fully available online (PROJECT_ID: Mouse_rest_psilocybin_pilot, https://openneuro.org/datasets/ds002154)
